# Divergence in bidirectional plant-soil feedbacks between montane annual and coastal perennial ecotypes of yellow monkeyflower (*Mimulus guttatus*)

**DOI:** 10.1101/2020.12.02.408245

**Authors:** Mariah M. McIntosh, Lorinda Bullington, Ylva Lekberg, Lila Fishman

**Affiliations:** Division of Biological Sciences, University of Montana, Missoula, Montana, USA; Department of Ecosystem and Conservation Sciences, University of Montana, Missoula, Montana, USA; MPG Ranch, Missoula, Montana, USA

**Keywords:** belowground ecology, life-history evolution, mycorrhizal fungi, soil ecology, plant–soil feedback, local adaptation

## Abstract

- Understanding the physiological and genetic mechanisms underlying plant variation in interactions with root-associated biota (RAB) requires a micro-evolutionary approach. We use locally adapted montane annual and coastal perennial ecotypes of *Mimulus guttatus* (yellow monkeyflower) to examine population-scale differences in plant-RAB-soil feedbacks.
- We characterized fungal communities for the two ecotypes *in-situ* and used a full-factorial greenhouse experiment to investigate the effects of plant ecotype, RAB source, and soil origin on plant performance and endophytic root fungal communities.
- The two ecotypes harbored different fungal communities and responsiveness to soil biota was highly context-dependent. Soil origin, RAB source, and plant ecotype all affected the intensity of biotic feedbacks on plant performance. Feedbacks were primarily negative, and we saw little evidence of local adaptation to either soils or RAB. Both RAB source and soil origin significantly shaped fungal communities in roots of experimental plants. Further, the perennial ecotype was more colonized by arbuscular mycorrhizal fungi (AMF) than the montane ecotype, and preferentially recruited home AMF taxa.
- Our results suggest life history divergence and distinct edaphic habitats shape plant responsiveness to RAB and influence specific associations with potentially mutualistic root endophytic fungi. Our results advance the mechanistic study of intraspecific variation in plant–soil–RAB interactions.

## INTRODUCTION

Plants cannot physically escape the abiotic and biotic properties of the soils where they germinate and are thus often adapted to local edaphic conditions. Plants may also alter soil properties and associated soil biotic communities with consequences for their own (and neighbors’) fitness, a process termed plant–soil feedback (PSF) (Bever *et al.*, 1997). PSF has become an important framework for understanding plant ecology and evolution, but dissecting the factors that shape plant–soil–microbe interactions remains a challenge (reviewed in Bennett & Klironomos, 2019). Plant functional traits, including life history, are predicted to shape plant responses to root-associated biota (RAB) (Berg & Smalla, 2009; Reinhart *et al.*, 2012; Koziol & Bever, 2015), as well as their effects on surrounding plant and soil microbial communities (Cortois *et al.*, 2016; Bardgett, 2017; reviewed in De Deyn, 2017). Understanding how specific plant traits interact with soil biotic communities (Reinhold-Hurek *et al.*, 2015; Cordovez *et al.*, 2019) may require a within-species perspective (Van Nuland *et al.*, 2016), but most studies of variation in PSF components have focused on differences among plant species (Kulmatiski *et al.*, 2008; Van Nuland *et al.*, 2016; Mariotte *et al.*, 2018). A micro-evolutionary approach may provide a window into the genetic and physiological mechanisms governing PSF interactions (Wilschut *et al.*, 2019; Mateus *et al.*, 2019; Pawlowski *et al.*, 2020).

Plant responses to soil biota can vary dramatically (Kulmatiski *et al.*, 2008; Reinhart *et al.*, 2012; Bennett & Klironomos, 2019), but most plant interactions with root-associated bacterial, fungal, and animal (e.g., nematode) communities are potentially harmful (Mandyam & Jumpponen, 2014). The fitness effects of plant-RAB interactions may depend on the evolutionary history and traits of each partner and the ecological context of soils and competitor species (Lekberg *et al.*, 2018; Bennett & Klironomos, 2019). Along with root traits (Semchenko *et al.*, 2018; Wilschut *et al.*, 2019), life history strategy may be particularly important in predicting belowground interactions (Koziol & Bever, 2015; Lemmermeyer *et al.*, 2015). Specifically, surveys have found that annual plants are often more negatively affected by the presence of native root-associated fungal communities (Reinhart *et al.*, 2012). Fast-growing ruderal plants (e.g., annuals or early-successional herbs) may generally suffer from interactions with soil biota. In contrast, long-lived perennials tend to experience fewer costs or even net benefits (Kulmatiski *et al.*, 2008). Such differences could involve multiple mechanisms, including both specificity of interactions (e.g., perennials preferentially interact with more mutualistic microbes and better exclude pathogens), as well as higher tolerance of perennials for similar colonization by pathogens or context-dependent mutualists (e.g., arbuscular mycorrhizal fungi; AMF). Late-successional species (which are often perennial) tend to be both more positively responsive to AMF than early-successional species and are more sensitive to fungal colonists (Koziol & Bever, 2015; Cheeke *et al.*, 2019).

Widespread or edaphically diverse plant species often encounter highly distinct soil biotas (Schechter & Bruns, 2008; Meadow & Zabinski, 2012), as climate and soil features structure biota across the landscape (Davison *et al.*, 2015; Bennett & Klironomos, 2019). Such spatial variation creates the potential for intraspecific adaptation to both biotic and abiotic belowground conditions. Indeed, a recent meta-analysis on local adaptation in plant interactions with mycorrhizal fungi found that plants generally performed better with all-home vs. away combinations of soil and AMF (Rúa *et al.*, 2016). Nonetheless, the generally negative net effect of live vs. sterile native soils (Kulmatiski *et al.*, 2008) suggests that plant adaptation to local soil biota may often not be sufficient to eliminate negative feedbacks. Furthermore, few studies have assessed factorial combinations of plant genotypes, soil biota, and soil in the same experiment (Rúa *et al.*, 2016), and thus cannot separately test for local adaptation to both biotic and abiotic components.

Along with variation in microbial feedbacks on plant performance, the PSF framework implies taxon-specificity in plant effects on root-associated biota. Indeed, while the RAB community largely reflects abiotic factors across broad spatial scales (Bennett & Klironomos, 2019), the microbes associated with a given plant species are generally not a random subset of those present in the local soil community (Hovatter *et al.*, 2011; Davison *et al.*, 2011; López-García *et al.*, 2014). This suggests selectivity in one or more of the partners and raises important questions about how it arises and is mediated (Baltrus, 2017). For example, studies of grasses have found heritable differences among plant genotypes in the rhizosphere bacterial community, despite temporal, soil, and climatic variability also affecting soil communities (Walters *et al.*, 2018). In contrast, a common garden study of the non-mycorrhizal model plant *Arabidopsis thaliana* found little effect of plant genotype on either eukaryotic or bacterial rhizosphere communities, despite local adaptation by the plants (Thiergart *et al.*, 2020). Questions of partner selectivity remain particularly open for endophytic, but putatively generalist, taxa such as AMF. Inoculation with the same AMF community can be strongly deleterious or positive for different plant species (Klironomos, 2003; Reinhart *et al.*, 2012) and environmental contexts (Larimer *et al.*, 2014), which may exert strong selection on plants to reduce costly associations and maximize beneficial ones. Further, AMF may compete with other soil biota for colonization of host taxa in root-limited environments. However, we are only beginning to understand the extent to which plant genotypes structure root-associated fungal communities.

Here, we leverage a fully factorial greenhouse experiment on yellow monkeyflower (*Mimulus guttatus,* Phrymaceae) ecotypes (Fig. 1) to investigate the interacting effects of plant genotype (including life history), soil origin, and RAB source (inoculum) on plant performance and root-associated fungal (RAF) communities. Yellow monkeyflowers are highly mycorrhizal (Bunn *et al.*, 2009; Meadow & Zabinski, 2012), exhibit extensive life history and edaphic diversity across a broad geographic range, and are a model system for evolutionary genomics. Thus, they provide an ideal species for investigating the mechanistic basis of feedbacks between plants and root-associated fungi. Our focal populations, a montane annual (Iron Mountain, Oregon; hereafter referred to as ‘IM’) and a coastal perennial (Florence Dunes, Oregon; hereafter referred to as ‘DUN’), represent widespread, genetically based life-history ecotypes adapted to ephemerally wet (annual) and summer wet (perennial) soils, respectively (Lowry & Willis, 2010). Thus, comparing these closely related ecotypes allows us to ask if patterns seen at large organizational levels (species, functional groups) are also evident at a micro-evolutionary scale.

**Figure 1.**
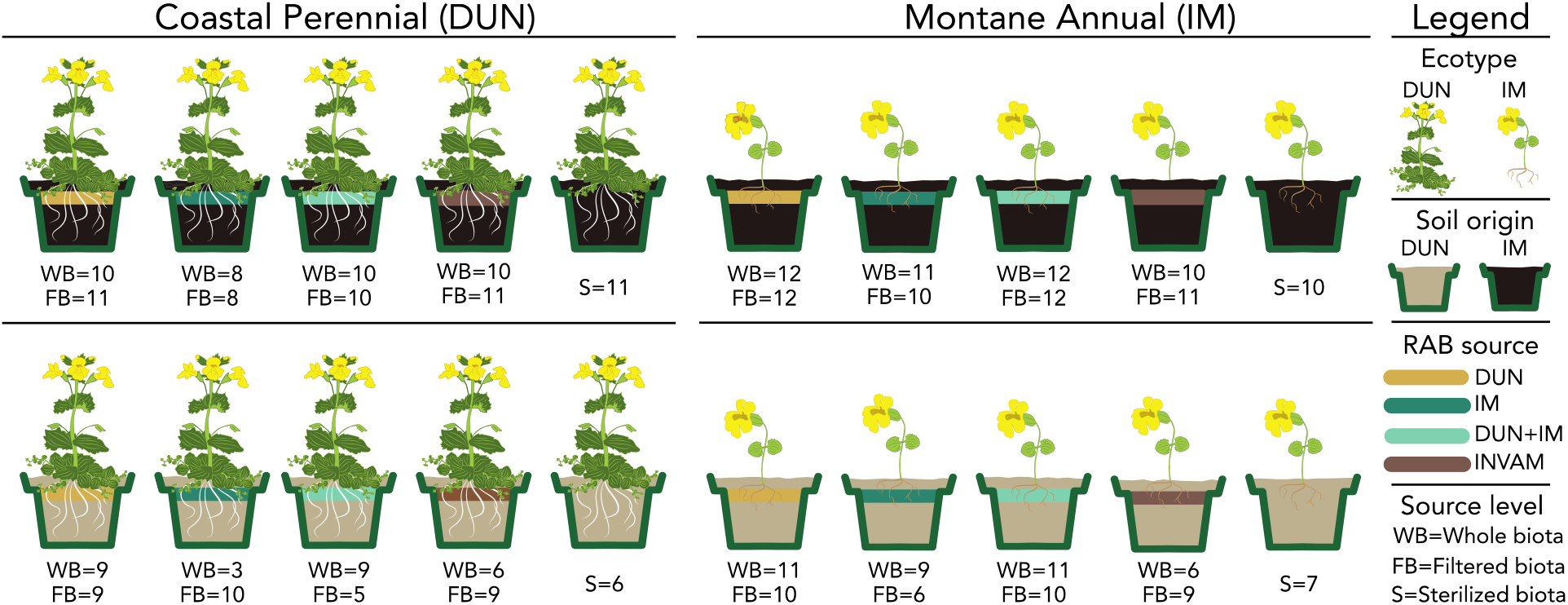
Design of common garden experiment examining feedbacks between *Mimulus guttatus* ecotypes and root-associated soil biota (RAB). Total N = 334. Sample size in each treatment after transplant mortality shown under each pot.

Our experimental design had three primary objectives, with the first two (*responsiveness* and *local adaptation*) focused on biotic and abiotic factors shaping plant performance. The third (*feedback*) focused on factors affecting the composition of root-associated fungal communities, which arguably include some of the most important pathogenic and mutualistic components of the soil and root biota (Bennett & Klironomos, 2019). After characterizing colonization of wild *M. guttatus* plants by fungal endophytes, we compared root and rhizosphere fungal communities within and among field sites using Illumina sequencing of the ITS2 region. We then generated population-specific soil biota as inocula and grew plants of each ecotype in a fully factorial greenhouse common garden design, including nine biotic treatments and two abiotic (sterilized) soil treatments (Fig. 1). We measured plant performance traits and compared performance and response intensity to evaluate the strength and direction of PSF for each ecotype under different conditions. We also explicitly tested for local adaptation by comparing ecotype performance in a greenhouse reciprocal transplant. Finally, we used Illumina sequencing to characterize the effects of plant ecotype, inoculum source, and soil on the composition of experimental RAF communities. Together, these analyses elucidate the biotic and abiotic factors underlying evolutionary shifts in PSFs within plant species.

## METHODS

### Study System

The *Mimulus guttatus* (Phrymaceae) species complex is noted for adaptation to extreme edaphic conditions, including copper mine tailings (MacNair, 1983), geothermal crusts (Lekberg *et al.*, 2012), and serpentine soils (Selby & Willis, 2018). A genetically based life-history polymorphism segregates across its broad latitudinal range in western North America (Alaska to Baja California). Annual populations grow in habitats with ephemeral water availability, while perennials grow in soils with year-round moisture (Lowry & Willis, 2010). The Iron Mountain (IM) montane annual population in the Oregon Cascades (1463 m elevation) consists of fast-growing non-rhizomatous plants that germinate in fall (overwintering as rosettes under snow) or after late spring snowmelt, and (if they survive to flower) often produce only a few flowers and fruits (Troth *et al.*, 2018). In contrast, the DUN perennial population (at sea level; Oregon Dunes National Recreation Area) experiences a moderate climate that is temperate year-round and receives continual moisture from rain, fog, or shallow pools in its rear-dune location (Hall *et al.*, 2006; Hall & Willis, 2006; Hall *et al.*, 2010). DUN exhibits a slower life history, spreading primarily through vegetative stolons, overwintering as rosettes, and flowering throughout the mild summer. IM soil is shallow, porous, and rocky, while DUN soil is little more than sand (Hall & Willis, 2006). At both sites, *M. guttatus* is a dominant member of the plant community during its growing season, although other (mostly perennial) herbs are also common at IM.

### Collection and characterization of field soils and RAF communities

#### Sampling of soils and roots

Paired samples of *Mimulus guttatus* roots and surrounding rhizosphere soils were collected at IM and DUN populations in June 2016 (n = 20 sampling sites per population). For the root samples, roots from multiple adjacent plants were collected on ice and pooled to obtain sufficient biomass per sample, and then split into subsets for sequencing analysis and staining for RAF colonization. The sequencing subsets were rinsed, then frozen at −80°C in 2mL tubes for later DNA extraction and library preparation (see below). The colonization subsets were dried in a drying oven for two days at 75 °C and stored in envelopes at room temperature until staining. Soil samples from the same sites were collected into 15mL tubes on ice and frozen at −80°C until DNA extraction. For soil nutrient analysis, a pooled soil sample from each site was analyzed by Ward Laboratories Inc. (Kearney, Nebraska, USA).

#### Evaluation of field RAF colonization

Twenty 2.5 cm long fine roots (~1mm diameter) per field sample were cleared in 10% KOH solution for 4 days, acidified in a 3% HCl solution overnight, rinsed, and then soaked in trypan blue dye solution for several days before being mounted and assessed under a microscope at 200X magnification. We scored AMF hyphae and vesicles and non-AMF hyphae (and sometimes microsclerotia or other non-AMF structures) as present or absent at 39–58 (mean = 44.8) intercepts per sample using the magnified intersection method (McGonigle *et al.*, 1990), and calculated proportion colonization for AMF and non-AMF structures by dividing the number of “present” intercepts by the total scored. AMF and non-AMF hyphae were separated by the former staining blue, lacking regular septa and having a more irregular shape and dichotomous branching pattern than non-AMF hyphae.

### Greenhouse experiment

#### Generation of experimental soils and inocula

In July 2015, we collected field soil from IM and DUN field sites to generate *M. guttatus*-associated inocula and for use as sterilized background soil in our greenhouse experiment. Soil from each site was collected within the rhizosphere of *M. guttatus* plants, pooled into plastic bins, and transported at ambient temperature to the University of Montana, where it was stored at 4°C. Pooling of field communities for experimental tests of plant responsiveness artificially reduces the within-site variance that plants experience in the wild (Reinhart & Rinella, 2016; but see Cahill *et al.*, 2017). However, inocula generated from the pooled field soils largely retained the broad community differences observed in the field root samples, which were properly replicated (see Results).

To generate background soil, bulk soil from each site was sterilized by autoclaving (three 99-minute cycles at 120 °C). From the live soils from each site, we generated paired whole-biota (WB) and filtered biota (FB) inocula (referred to hereafter as RAB levels) from DUN and IM root/soil samples, using native plants and soils under greenhouse conditions. We also generated a 1:1 DUN + IM mixed inoculum (both WB and FB levels) and an AMF-enriched inoculum originally obtained from the INVAM collection. All inocula were used to inoculate experimental treatments (See Supporting information Methods S1 for details on inocula generation).

#### Plant growth conditions

To set up factorial treatments (Fig. 1), 200mL Cone-Tainer pots were filled with 100mL of a mix (1:1:1) of sterilized field soil (DUN or IM), Turface, and sterile sand, inoculated with 30mL of the appropriate inoculum (WB and FB for each of the four sources) or sterilized field soil (S), and topped off with the appropriate soil mixture. Previously collected *M. guttatus* seeds from DUN and IM populations were cold treated for one week (4°C) on wet sterile sand in sealed petri dishes, germinated at greenhouse temperatures, and transplanted into the treatment pots. If necessary, we re-transplanted up to three times within ten days of the original transplanting to obtain one plant per pot. Pots were randomly arrayed in stands to avoid greenhouse effects but blocked (in sets of eight) by inoculum in order to avoid cross-contamination. Plants grew under (flowering-inductive) 16-hour days, which mimicked summer field conditions, with daily or twice-daily watering to maintain saturation and biweekly (first 2 months) and then weekly fertilization with 10mL of 20-2-20 fertilizer at 50ppm N.

#### Performance measures and sample collection

Plants grew until the majority had flowered and were harvested before they began to senesce (approximately 11 weeks). Prior to harvest, we recorded the date of the first flower (days from transplant date) and the total number of flowers. A few plants that did not open any flowers before harvest (but generally had buds) were assigned a flowering time of 75 days (3 days after the maximum recorded) and a flower count of 0; this provides a more accurate record of plant performance than treating those values as missing data. We separated and dried belowground and aboveground biomass, subsampling roots for colonization estimates and molecular analysis into 96-well plates on ice, storing them at −80°C until use. We dried biomass samples in a drying oven at 70 °C until dry and then recorded dry weight.

#### Statistical analyses of plant performance

To address the effect of plant ecotype, soil, and soil biota completeness on plant performance, we first used a fully factorial ANOVA with ecotype, soil and RAB level—whole soil (WB), filtered (FB), and sterilized (S) treatments—as factors, combining all RAB sources. To examine context-dependent responsiveness to the different whole soil biota (RAB source; DUN, IM, DUN+IM, INVAM) we also calculated the Relative Interaction Intensity (RII; following Waller *et al.*, 2016) for each WB performance value, standardized by the mean value for that ecotype in that soil (RII = (Performance_WholeSoil_E, S_ – mean Performance_Sterile_E, S_,)/ Performance_WholeSoil_E, S_ + mean Performance_Sterile_E, S_, where E and S are ecotypes and soils). RII (which is bounded by −1 and 1) scales plant performance in whole soil biota relative to the matched sterile treatment, allowing comparisons of responsiveness despite major ecotype and soil effects on absolute trait values. We then conducted a full factorial ANOVA (with soil, ecotype and RAB source as factors) on RII values for each trait. To characterize differences in the strength and nature of joint biotic and abiotic soil feedbacks between DUN and IM populations, we also explicitly compared RII in home combinations, using an ANOVA with site (matched ecotype, soil, and source), RAB level (WB vs. FB), and their interaction. Finally, to assess local adaptation to belowground environments, we compared plant performance in all-home (matched ecotype, soil, and source) vs. away (unmatched ecotype, soil, and source) for the subset of treatments with DUN and IM WB inoculum. All ANOVA and Tukey’s Honest Significant Difference post-hoc comparisons of means were implemented JMP 14.0 (SAS Institute, 2018).

### Characterization of experimental colonization and root-associated fungal communities

We scored a sample of experimental roots (N = 3 per treatment) for colonization by AMF structures in following the protocols described above for field roots. In addition, we scored root intersects for non-AMF structures, including hyphae and other structures from dark septate endophytes (Jumpponen & Trappe, 1998).

We extracted DNA from field root samples (freeze dried) using PowerLyzer PowerSoil DNA Isolation Kits (MoBio, now Qiagen; Hilden, Germany) and from fresh greenhouse roots using a 96-well format CTAB-chloroform protocol (dx.doi.org/10.17504/protocols.io.bgv6jw9e). We used a two-part PCR protocol (Supporting Information Table S1) to amplify and barcode the fungal ITS2 region. We sequenced all samples on Illumina MiSeq platforms (2 x 300 bp). For more details on sample preparation, amplification, and sequencing, see Supporting Information Methods S2.

We used ‘Quantitative Insights Into Microbial Ecology 2’ (Bolyen *et al.*, 2019) for initial bioinformatics analysis of RAF sequences (for more details, see Supporting Information Methods S3). To assign fungal taxa to functional categories, we used FUNGuild (Nguyen *et al.*, 2016) as described in Supporting Information Methods S3. Low taxonomic and ecological resolution of knowledge about fungal sequence variants limits inferences from these categorical assignments, but they capture major functional differences in fungal biota among sites and treatments (Nguyen *et al.*, 2016).

Fungal communities were characterized and compared using the VEGAN package (Oksanen *et al.*, 2017) implemented in R (R Core Team 2018, v 3.5.1) and R Studio version 1.1.453, unless otherwise noted. We rarefied the field and greenhouse sequence datasets at 1700 and 200 sequences per sample, respectively (see Supporting Information Fig S1 for rarefaction curves). We performed non-metric multidimensional scaling (NMDS) on Bray-Curtis distances of Hellinger-transformed sequence abundances to compare community composition among field sites as well as treatments in the greenhouse. Each NMDS analysis was performed using the metaMDS function in VEGAN. The adonis function for permutational multivariate analysis of variance (PERMANOVA) was used to test for significant variation in community composition recovered, using 1000 permutations (Oksanen *et al.*, 2017). Differences in overall richness as well as proportions of functional groups among field sites and within the greenhouse treatments were assessed using a two-way ANOVA and Tukey’s HSD for pairwise comparisons. To explore the contributors to WB community differences, we used one-way ANOVAs to test taxa with >10 reads (across the entire dataset) for differences in abundance by ecotype, soil and RAB source (with false discovery rate protection for multiple tests in JMP14; SAS Institute, 2018).

## RESULTS

### Field soil and root fungal communities and soil-to-root filtering differ between coastal and montane *M. guttatus* populations

Abiotic soil properties differed substantially between coastal perennial (DUN) and montane annual (IM) *Mimulus guttatus* populations, with IM soil substantially (15-20x) higher in organic matter, nitrogen, and magnesium than DUN soil (Supporting Information Table S2). Consistent with these abiotic differences, RAF community richness (sequence variants) in IM rhizosphere soil was twice as high as in DUN soil (richness = 206 ± 10 vs. 97 ± 10, p<0.01). Site was also the most important factor structuring RAF communities (Fig. 2A), although medium (soil vs. root) and a site*medium interaction were also highly significant (Supporting Information Table S3). FUNGuild analysis indicated substantial filtering by *M. guttatus* roots relative to their surrounding soil communities; DUN root samples were significantly enriched for mutualists, whereas IM roots were enriched for pathogens (Fig. 2B). The most common AMF sequence variants at DUN were *Claroideoglomus* (Claroideoglomeracae; mean reads/sample = 23 ± 10), while the only putative mutualists substantially present in IM roots (mean > 2 reads/sample) were assigned to family Glomeraceae (mean reads = 19 ± 17). Nonetheless, DUN and IM roots exhibited similarly low colonization by AMF (7-12%) and non-AMF endophytic structures (45-54%; Supporting Information Table S4). Strong differences between IM and DUN RAF communities were maintained in their respective inocula (Fig. 2A); in particular, *Claroideoglomus* and Glomeraceae remained the dominant AMF taxa in DUN and IM inoculum samples, respectively.

**Figure 2.**
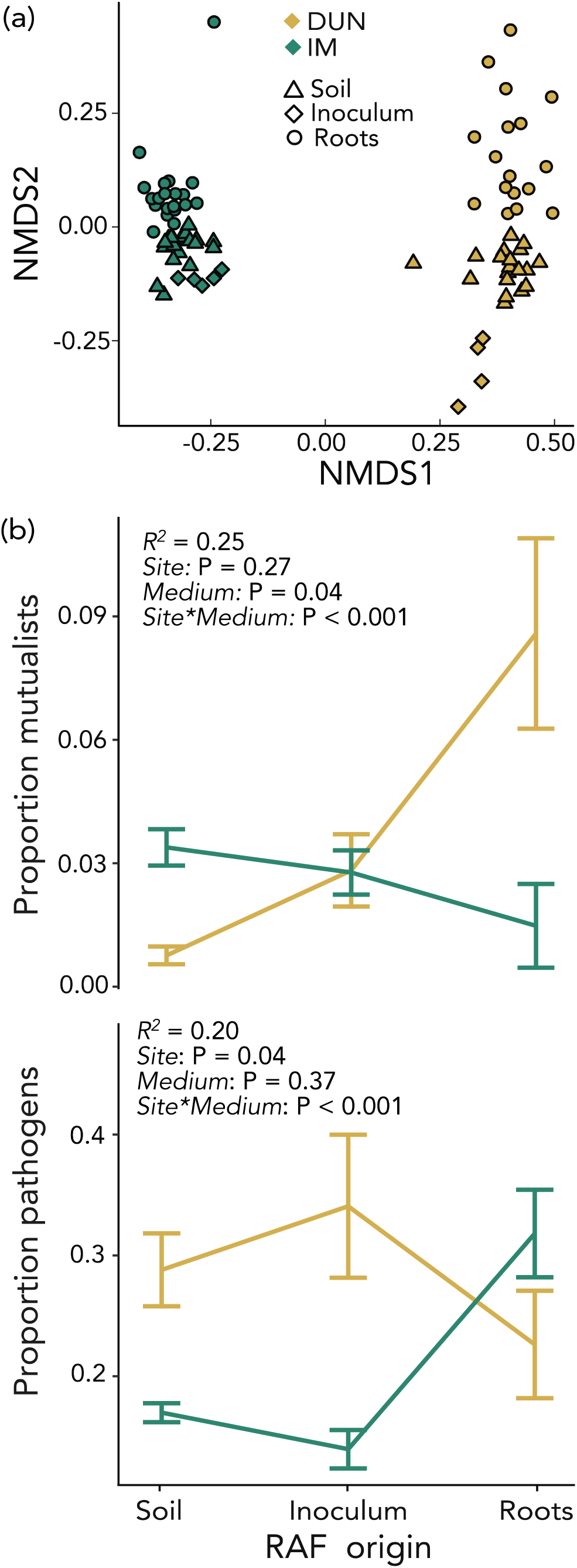
Root-associated fungal (RAF) communities at coastal perennial (DUN) and montane annual (IM) *Mimulus guttatus* field sites. A) Non-metric multidimensional scaling (NMDS) on Bray-Curtis distances of Hellinger-transformed sequence abundances. Colors correspond to site (DUN = tan, IM = green), while shapes correspond to fungal communities sampled (triangle = rhizosphere soil, diamond = inoculum, circle = roots). B) Mean (± SE) proportion of sequences categorized by FUNGuild as probable mutualists and pathogens at each site and sample category.

### Plant performance depends on ecotype, soil and their interactions, but also on soil biota

In the common garden, ecotypes were significantly differentiated for all traits except aboveground biomass. Soil origin (DUN, IM) affected all plant performance metrics, and RAB level (WB, FB, S) influenced aboveground and total biomass as well as flower number (P <0.005 in full factorial models, Table 1). DUN plants flowered later and made fewer flowers than IM plants but allocated more to belowground biomass and were larger overall (Table 1, Fig. 3). Plants grown in DUN soil, which is nutrient-poor and mostly sand (leading to visibly lower water retention during the experiment, L. Fishman, pers. obs.), performed poorly vs. those in IM soil (Fig. 3). However, strong soil*ecotype interactions for flower number and belowground biomass resulted from high plasticity for these traits in IM and DUN plants, respectively (Table 1, Fig. 3). In the richer IM soil, each ecotype allocated more to the trait potentially important for fitness for its life history (e.g., flowers in annual IM, belowground biomass in perennial DUN). Performance for several traits was lower in whole soil (WB) vs. sterilized (S) treatments (Fig. 3, Table 1), and WB plants also had significantly lower aboveground and total biomass than FB (filtered biota) plants (Tukey’s HSD *P* < 0.05). This suggests that fungal endophytes (which were largely removed by filtration; Supporting Information Methods 1) contributed substantially to the generally negative biotic feedbacks. Biotic feedbacks also interacted with soils for aboveground and total biomass (*P* < 0.05 for soil*RAB source interaction), with the nutrient-rich IM soil exacerbating the negative effects of complete WB biota (Fig. 3, Table 1).

**Table 1.** Model analysis for performance traits across *Mimulus guttatus* ecotype (DUN, IM), soil (DUN, IM), and broad biotic levels in root-associated soil biota (whole biota, filtered biota, sterilized). F-ratios from full factorial ANOVA model (Poisson GLM with log-link for flower number) are highlighted by their statistical significance **(bold**: < 0.0001, ***bold italic***: <0.005, *italic* < 0.05).

| Factor | df | Days to Flower | Flower number | AG Biomass | BG Biomass | Total Biomass |
| --- | --- | --- | --- | --- | --- | --- |
| Ecotype | 1 | **162.38** | **172.32** | 3.42 | **135.83** | **54.70** |
| Soil | 1 | **13.40** | **47.56** | **196.08** | **74.18** | **179.72** |
| Level | 2 | 3.03 | **6.00** | **21.75** | 1.16 | **11.47** |
| Ecotype*Soil | 1 | 0.34 | **10.11** | 0.39 | **13.53** | 2.39 |
| Ecotype*Level | 2 | 1.11 | 1.90 | 1.38 | 1.59 | 1.13 |
| Soil*Level | 2 | 1.99 | 2.97 | **5.84** | 0.54 | 3.20 |
| Ecotype*Soil* Level | 2 | 0.16 | 0.93 | 1.16 | 1.64 | 0.74 |

**Figure 3.**
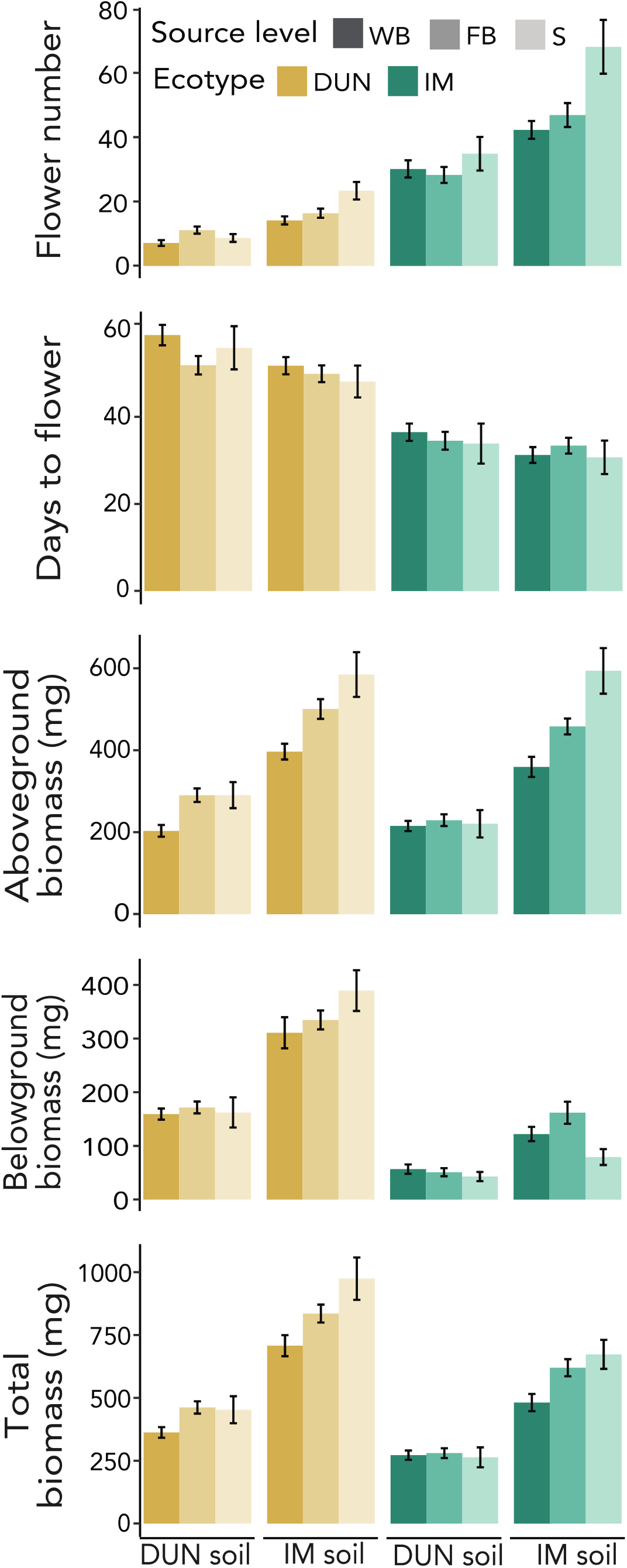
Mean (± SE) performance of each *M. guttatus* ecotype (DUN, IM) grown in each soil (DUN, IM) and with whole biota (WB), filtered biota (FB), and sterilized (S) RAB levels (across all RAB sources).

### Plant responsiveness to particular soil biota varies across performance traits, ecotypes, soils, and RAB sources, but reveals strong negative biotic feedbacks rather than local adaptation

For examining plant responses to distinct RAB source communities, we focused on the WB treatments, using Relative Interaction Intensity (RII; see Methods) to control for the main effects of ecotype and soil. RII varied substantially across the four RAB sources (DUN, IM, DUN+IM, INVAM; *P* < 0.01 for all traits). Plants were generally less negatively (or positively for belowground biomass) affected by the AMF-rich INVAM RAB and the DUN RAB compared to the IM and DUN+IM RAB (Table 2, Fig. 4). Across soils and RAB, ecotypes differed significantly in RII for belowground biomass (*P* < 0.001), with IM tending to respond more positively. However, significant ecotype*soil (aboveground and belowground biomass) and ecotype*RAB source (flower number) interactions (Table 2) resulted from weak RII for biomass traits exhibited by IM plants in DUN relative to IM soil (Fig. 4).

**Table 2.** Model analysis for relative interaction intensity (RII) across *Mimulus guttatus* ecotypes (DUN, IM), soils (DUN, IM), and whole biota RAB sources (DUN, IM, DUN+IM, INVAM). RII standardizes performance relative to the same ecotype and soil combination without biotic inoculation (sterilized), so ecotype and soil effects indicate variation in responsiveness to biota per se. F-ratios from full factorial ANOVA model are highlighted by their statistical significance (**bold**: < 0.0001, ***bold italic***: <0.005, *italic* < 0.05).

| Factor | df | Days to Flower | Flower number | AG Biomass | BG Biomass | Total Biomass |
| --- | --- | --- | --- | --- | --- | --- |
| Ecotype | 1 | 0.14 | 0.55 | 3.60 | **8.50** | 3.21 |
| Soil | 1 | 0.59 | **10.10** | **21.03** | 0.13 | **16.52** |
| Source | 3 | *3.51* | **5.77** | **18.61** | **5.58** | **14.64** |
| Ecotype*Soil | 1 | 1.74 | 1.06 | **14.88** | 5.56 | 3.72 |
| Ecotype*Source | 3 | 0.87 | 4.30 | 1.26 | 0.02 | 0.71 |
| Soil*Source | 3 | 0.85 | 2.71 | 1.55 | 0.71 | 2.18 |
| Ecotype*Soil* Source | 3 | 1.17 | 3.17 | 1.62 | 3.64 | 1.62 |

**Figure 4.**
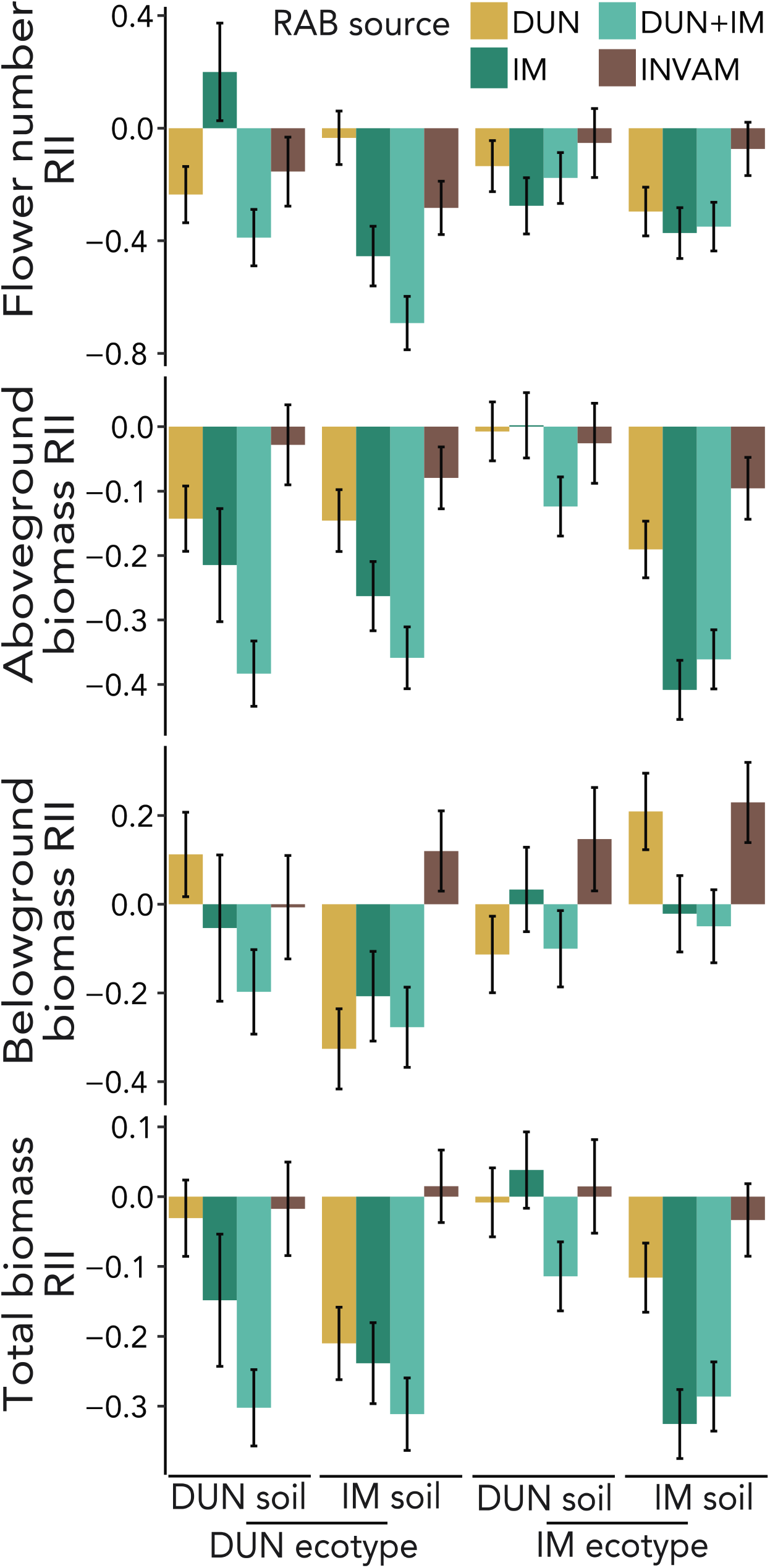
Responsiveness (RII; mean ± SE) of performance traits to four sources of whole soil RAB for DUN and IM *M. guttatus* plants grown in DUN and IM soil. RII standardizes individual performance in a biotic treatment by the mean for that same ecotype and soil, removing the direct effects of those factors on the traits.

To assess local adaptation, we compared the aboveground biomass (a common proxy for fitness that does not vary among our common garden-grown ecotypes) of the ecotypes in home and away sterile soil, as well as all-home vs. all-away (soil and WB biota RAB source matched to site) combinations. There was no interaction effect of soil alone with ecotype (*P* = 0.57), but a near-significant ecotype*site interaction (*P* = 0.12; Supporting Information Table S5) resulting from *lower* performance (population-specific negative feedback) rather than enhanced performance (local adaptation) in all-home vs. all-away soil conditions (i.e., a virtual reciprocal transplant). To unpack this further, we directly compared RII (which accounts for soil and ecotype differences) for DUN and IM in their respective home FB and WB biota. Both aboveground and total biomass exhibited strong site*RAB level interaction effects; IM was strongly negatively responsive to home WB biota (but not the home FB biota) in home soils, while DUN was relatively unaffected (RII essentially zero) to the presence of both home FB and WB biota in home soils (Fig. 5). Thus, IM soil conditions appear particularly conducive to negative biotic feedbacks from native fungi, and IM plants appear vulnerable to those local biotic feedbacks.

**Figure 5.**
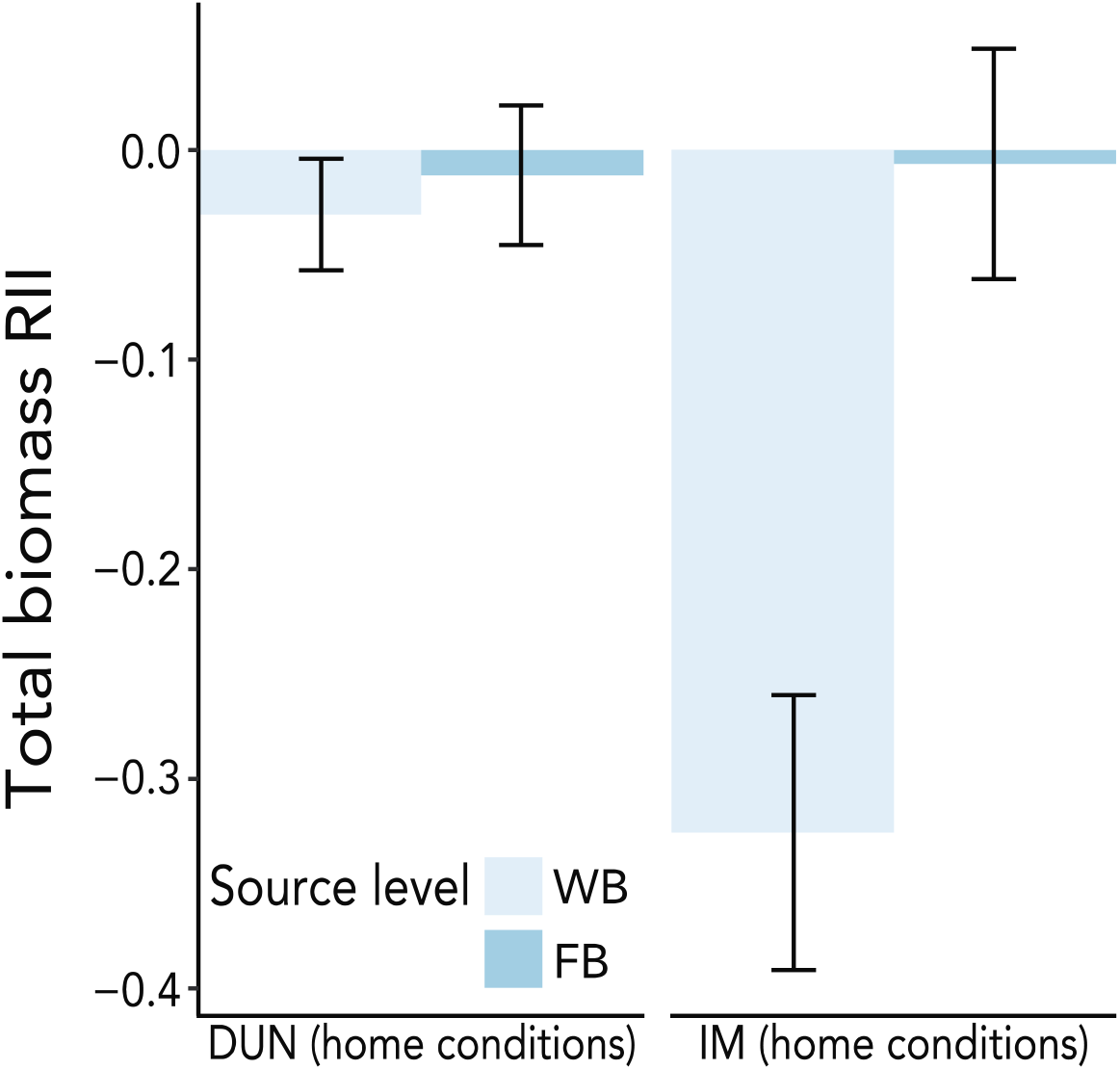
Biomass feedbacks (RII mean ± SE) on *M. guttatus* ecotypes of home filtered biota (FB) and whole biota (WB)home soils. Higher negative responsiveness of IM annuals of interactions with its home endophyte-containing (WB) community is the opposite of the pattern expected from local adaptation to soil biota.

### Plant ecotype, soil, and RAB source affect root colonization by fungal endophytes

In WB samples, soil and RAB source strongly influenced both AMF and non-AMF colonization (all *P* < 0.005 in ANOVA; Table 3). Root colonization by AMF was higher in IM vs. DUN soil (0.21 ± 0.02 and 0.10 ± 0.02, respectively), while non-AMF colonization showed the opposite pattern (0.27 ± 0.04 vs. 0.33 ± 0.03, respectively). In terms of RAB source, root colonization by AMF was high in plants inoculated with INVAM (0.23 ± 0.04) and DUN+IM (0.21 ± 0.03) inocula, and lower in the pure DUN (0.07 ± 0.03) and IM (0.10 ± 0.03) inocula. High AMF root colonization in the INVAM inoculum suggests the relatively benign effects of this RAB (Fig. 3) do not result from *Mimulus* not being colonized in this treatment. Non-AMF colonization was notably high in treatments containing IM biota (IM: 0.35 ± 0.05, DUN+IM: 0.46± 0.05; *P* <0.001 in LSMs contrast vs. others), and microsclerotia were common in these samples relative to DUN-inoculated roots (M. McIntosh and Y. Lekberg, pers. obs.). Small sample sizes for the three-way combinations limit direct statistical comparison of home-DUN to home-IM colonization patterns. However, high non-AMF colonization in IM WB inocula may contribute to the low performance of IM plants under home conditions (Fig. 5). Plant ecotype also significantly influenced AMF root colonization (*P* = 0.03), but not non-AMF colonization (*P* = 0.28) in the full model, with the perennial DUN roots nearly twice as colonized by AMF as IM roots across all treatments (LSMs: 0.19 vs. 0.12 ± 0.02, N = 25 and 17, respectively; Table 3). Ecotypic differences in AM colonization suggest that the plants are differentiated in interactions with fungi regardless of local abiotic or biotic soil conditions.

**Table 3.** AMF and non-AMF colonization of *M. guttatus* roots differs between DUN and IM ecotypes and varies among experimental soil (DUN, IM) and wild-derived RAB sources (DUN, IM, DUN+IM) treatments.

| Source | df | AMF | AMF | Non-AMF | Non-AMF |
| --- | --- | --- | --- | --- | --- |
|  |  | F Ratio | Prob > F | F Ratio | Prob > F |
| Soil | 1 | 10.705 | 0.0028* | 10.999 | 0.0025* |
| Ecotype | 1 | 4.711 | 0.038* | 1.206 | 0.28 |
| RAB Source | 3 | 5.369 | 0.0046* | 11.115 | <.0001* |
| Soil*RAB Source | 3 | 4.176 | 0.0142* | 0.265 | 0.61 |
| Ecotype*RAB Source | 3 | 2.508 | 0.08 | 2.488 | 0.08 |
| Soil*Ecotype | 1 | 1.654 | 0.21 | 0.851 | 0.48 |

### Experimental fungal communities are broadly defined by source inoculum and soil, but the AMF community also depends on interactions with plant ecotype

In the greenhouse common garden, the composition of root-associated fungal (RAF) communities in the three *Mimulus*-derived WB treatments was explained primarily by RAB source (inoculum)(r^2^ = 0.15, *P* < 0.01 in PERMANOVA). Four taxa with highly significant differences among ecotypes generated this pattern (all FDR-protected *P* < 0.005, no others with *P* <0.05): the dark septate endophyte *Alternaria* and the AMF family Glomeraceae (both common in IM, rare in DUN, intermediate in DUN+IM), the AMF genus *Claroideoglomus*, and a functionally uncategorized *Sebacinale* (both common in DUN and DUN+IM, rare in IM) (Supporting Information Table S6). The particularly high abundance of *Alternaria* in DUN+IM and IM inoculated roots (LSMs contrast: *P* < 0.0001), and IM soil (*P* = 0.04) in the greenhouse experiment, along with abundant non-AMF structures in these treatments, makes it a strong candidate for explaining relatively poor plant performance in both IM-including inocula (Fig. 4) and IM home conditions (Fig. 5). Site-specificity of strong negative interactions with DSE may also occur in the field, as *Alternaria* abundance was elevated in roots vs. soil at IM (74.0 ± 12.1 SE and 23.7 ± 12.1 SE; site*medium interaction *P* = 0.04), but constant at DUN (32.4 ± 13.6 SE and 34 ± 12.5 SE, respectively).

Overall, whole RAF communities did not differ between DUN and IM ecotypes (*P* = 0.52), but they did marginally between soil origin (r^2^ = 0.04, *P* < 0.05). Because two of the most site-differentiated taxa in both the field and inoculated roots were AMF, however, we also specifically examined shifts in the AMF community among ecotypes. Despite the strong effect of RAB source, AMF in roots exhibited an interesting ecotype*source interaction (Fig. 6, Supplemental Information Table S6). *Claroideoglomus* predominated in roots of both ecotypes in DUN and DUN+IM RAB sources (proportion *Claroideoglomus* > 95%), and IM plants were exclusively colonized by Glomeraceae AMF in the IM inoculum, but five of nine DUN plants grown in the Glomeraceae-rich IM inoculum fostered *Claroideoglomus* (mean proportion *Claroideoglomus* = 23%), producing the interaction in abundance. Thus, the coastal perennial ecotype appears to preferentially recruit rare “home” AMF taxa present in the “away” RAB.

**Figure 6.**
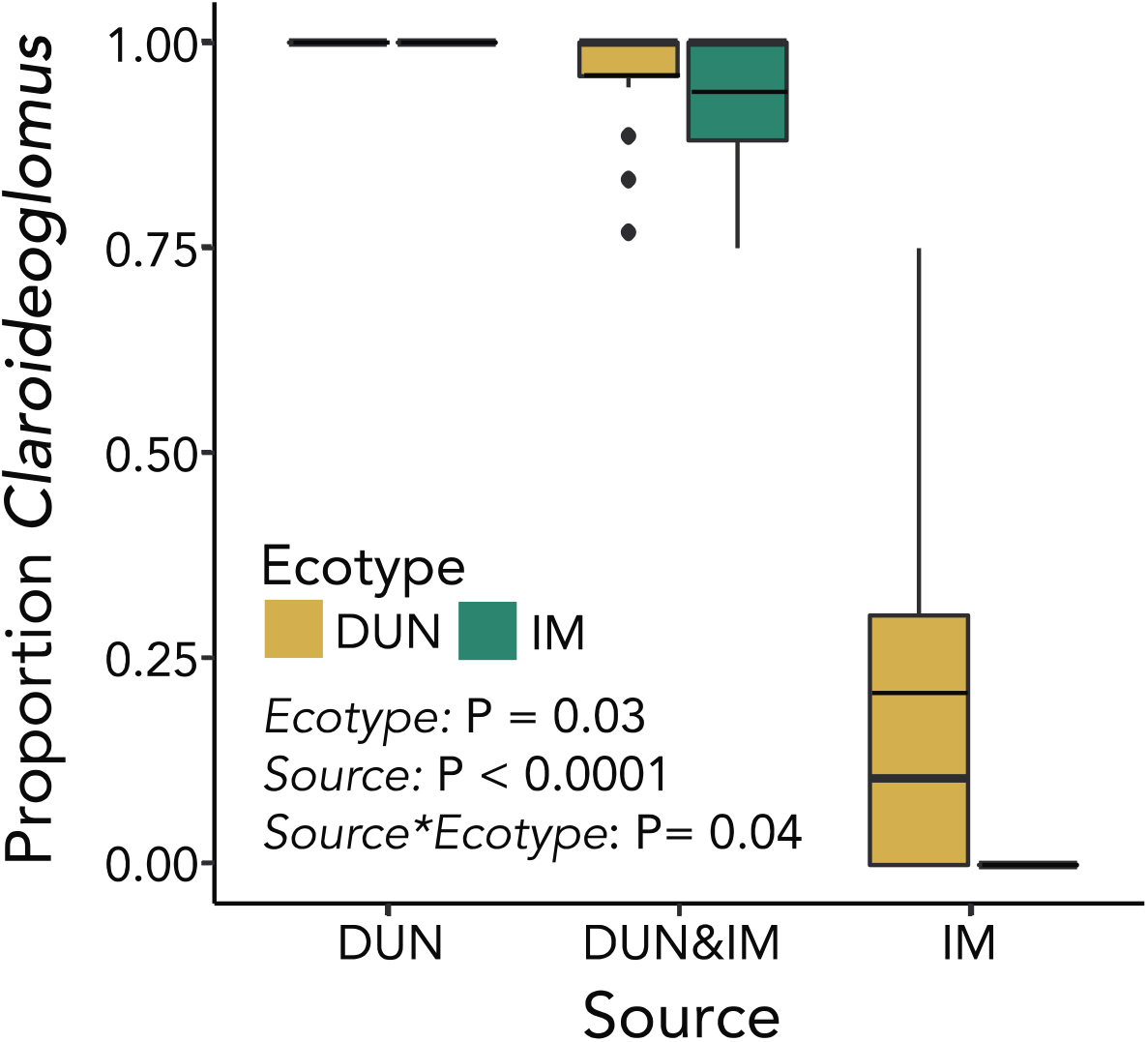
Abundance of DUN-dominant AMF taxon *Claroideoglomus* (as a proportion of total AMF reads) in roots of IM and DUN *M. guttatus* grown in soils inoculated with DUN, DUN+IM, and IM whole biota. Total N = 54. Bold lines across boxes represent the median of each group, while thin lines show the mean. The whiskers extend ≤1.5*IQR, while outlier beyond 1.5*IQR are plotted individually.

## DISCUSSION

Understanding how plant and microbial traits shape the outcomes of below-ground interactions (and vice versa) remains a key challenge in ecology and, increasingly, evolutionary biology. Differences in plant-soil feedbacks (PSF) among plant species (e.g. Bennett & Klironomos, 2019) arise from divergence among populations; thus, an intraspecific perspective may illuminate the evolutionary origins of PSF components. Here, we used representatives of widespread *Mimulus guttatus* ecotypes to ask whether and how intraspecific differences in habitat and life history (along with abiotic and biotic soil conditions) influence the magnitude and direction of feedbacks on reproductive and vegetative performance traits, as well as plant responsiveness to microbial communities. We tested for local adaptation by plants to distinct microbial/soil conditions and examined how soils and plants shape root-associated fungal communities over the short term. In addition to revealing the complexity of PSF from both plant and fungal perspectives, our results provide a window into the early stages of divergence in PSF among plant populations, an important step toward understanding its mechanistic basis and a key consideration for management of plant populations.

### Life history and edaphic differentiation in yellow monkeyflowers generates distinct contexts for interactions with root-associated fungi (RAF)

As expected for sites with distinct edaphic and climatic conditions (Tedersoo *et al.*, 2014), RAF were strongly differentiated between montane and coastal *Mimulus guttatus* population *s.* Higher soil nutrients (Supporting Information Table S2) and plant cover at the montane IM site likely generate more niches for all RAF (Waldrop *et al.*, 2006; Tedersoo *et al.*, 2014), explaining its higher richness vs. the sandy and flooded swale habitat at DUN. *Mimulus* roots substantially filtered the RAF community in the field, as is generally observed (Hempel *et al.*, 2007; Goldmann *et al.*, 2016), with DUN root samples enriched for endophytic mutualists (primarily AMF) and IM root samples enriched for pathogenic taxa relative to surrounding soils (Fig. 2). Roots from both sites were colonized by AMF at similar levels, confirming that yellow monkeyflowers are mycorrhizal (Bunn & Zabinski, 2003; Meadow & Zabinski, 2012). However, the AMF taxa represented in roots were largely non-overlapping at the family level, with Claroideoglomeracae most common at DUN, but Glomeraceae dominant at IM. Tolerance of low nutrient conditions may explain the dominance of *Claroideoglomus* in the sandy DUN soil, while the absence of Gigasporaceae there breaks the broad association of this large-spored group with maritime soils (Stürmer *et al.*, 2018). Even moderate colonization of field *M. guttatus* roots by AMF (primarily Glomeracae) at the annual IM site is notable, as the summer growing season often lasts only a few weeks before plants die of drought (Troth *et al.*, 2018; Nelson *et al.*, 2018). However, because most plants at IM germinate in the fall (Nelson *et al.*, 2018), initial AMF colonization may occur before rosettes overwinter. Nonetheless, the short growing season at IM may contribute to its guild-level filtering toward pathogens, as *Mimulus guttatus* may be a key year-round carbon-source for AMF at the low-cover DUN site but a relatively ephemeral resource for mutualists at IM. The field differences in RAF also create the opportunity for, and may in part reflect, evolved differences between the *Mimulus guttatus* ecotypes in their interactions with fungal communities (see below).

### Negative feedbacks on plant performance are highly context-dependent, but ecotypic differences in responsiveness reflect life history divergence rather than local adaptation

In our factorial experiment, plant ecotype, soil, and both the level and specific source of soil biota (as well as interactions) affected aspects of plant performance (Figs. 3-5). Overall, under short-term greenhouse growth conditions, the sum of soil biotic feedbacks on yellow monkeyflowers was primarily negative (Figs. 3, 4), as is often seen in herbaceous plants (Kardol *et al.*, 2006; Kulmatiski *et al.*, 2008; Lekberg *et al.*, 2018). Further, the generally worse effects of whole soil vs. filtered (large propagules removed) inocula (Fig. 3) suggests that AMF do not mitigate negative effects of other components of the community or may be parasitic themselves. This is expected under conditions where carbon costs of association are not compensated for by nutrient or stress-tolerance benefits (Werner & Kiers, 2015). However, the INVAM inoculum had relatively weak (or no) negative feedbacks on plant performance (Fig. 4), supporting the idea that generic AMF inoculations dampen differences among plant ecotypes in responsiveness compared to native whole soil biota (Rúa *et al.*, 2016). Overall, our joint performance, colonization and community data further illustrate the complexity of PSF interactions now being revealed in many systems (Bennett & Klironomos, 2019), and emphasize that net effects on plant performance integrate across potentially conflicting abiotic and biotic factors.

Relative Interaction Intensity (RII) also varied strongly among combinations of soil, RAB source, and ecotype for all traits except flowering time. Genetic differentiation within *M. guttatus* in the magnitude and nature of feedbacks from soil biota was evident in different contexts, with DUN experiencing overall stronger negative feedbacks on belowground biomass and, to a lesser extent, flower number and aboveground biomass (Fig. 4). Community-level studies have suggested that late-successional species and perennials are more positively AMF-responsive and have higher specificity (Reinhart *et al.*, 2012; Cheeke *et al.*, 2019). Given the higher AMF colonization rates in DUN-source RAB and evidence for specificity of AMF interactions (see below), net negative RII for these traits in DUN may result from intense interactions with root endophytes that are adaptive during multi-year growth in native settings but deleterious in a short-term (and higher nutrient) pot experiment.

To isolate the belowground interactions that plants experience in the field and test for local adaptation, we also asked (as most studies of PSF across species do; Kulmatiski *et al.*, 2008) how each ecotype responded to the presence or absence of home biota in home soils. In this home-only comparison, IM annuals experience much stronger negative biotic feedbacks than DUN perennials for aboveground and total biomass (Table 2, Figs. 4, 5).

Although plant-soil feedbacks affected most traits, plants in all-home (vs. all-away) soil+biota combinations did not exhibit the relatively high-performance characteristic of local adaptation. Thus, the coastal-montane adaptation evident in *M. guttatus* reciprocal transplants (Hall & Willis, 2006; Popovic & Lowry, 2020) may be mediated primarily by climate or other factors. Local adaptation appears common in home vs. away plant-soil-AMF combinations (Rúa *et al.*, 2016), but non-AMF negative interactions in our whole soil treatments may outweigh any locally-adapted AMF associations (see below). Further, relatively low-stress greenhouse conditions (e.g., higher N) may decrease both the opportunity for and expression of local adaptation by both microbes and plants (Johnson *et al.*, 2010) and may also exacerbate pathogen abundance and disease (Walters & Bingham, 2007). Finally, the distinct life history strategies of our ecotypes may foster local adaptation to tolerate or encourage high colonization by fungal endophytes at DUN (as evidenced by minimal biotic feedbacks under native conditions; Fig 5). Conversely, strong negative WB feedbacks on the annual under home conditions (Fig. 5) may be a byproduct of a life history strategy evolved to rapidly capitalize on ephemeral resources for reproduction at the expense of defense against pathogens. Aboveground anti-herbivore defense in *M. guttatus* is structured (in part) by life history and flowering time, with high elevation annuals like IM among the least defended (Kooyers *et al.*, 2017). High vulnerability to native fungal biota in this experiment (which lasted longer than a typical IM growing season) suggests a similar belowground avoidance (vs. resistance/tolerance) strategy in IM annuals.

While our results recapitulate several expected patterns (e.g., generally negative feedbacks of whole soil communities on plant performance), the numerous quantitative interactions underline the complexity in interpreting broad analyses of PSF and plant performance of many species across habitats (Kulmatiski *et al.*, 2008; Cortois *et al.*, 2016). Through differences in root architecture and timing of growth, annuals and perennials may interact with distinct soils and biota even at the same site, as well as express intrinsic differences in responsiveness to the same soil fungi. Our factorial approach indicates that all of these factors may matter for the strength of PSF, even when comparing just two populations of the same plant species. As it becomes possible to more finely characterize the functional characteristics of root-associated biota (Agler *et al.*, 2016), this picture is likely to become even more complex.

### Ecotypic differences in endophyte colonization and specificity of AMF associations suggest intraspecific genetic divergence in plant-fungal interactions

Our factorial experiment, combined with RAF sequencing, can begin to tease apart how plant traits associated with *M. guttatus* ecotype shape RAF communities, and vice versa. Experimental root fungal communities were primarily inherited from their source inoculum, mirroring the biogeographic factors that shaped the source RAF (Hoeksema *et al.*, 2010; Bennett & Klironomos, 2019). In addition, whole-community sequencing reveals that non-AMF endophytes, such as DSE, likely play an important role in determining plant performance. Most notably, the two treatments (IM and DUN+IM inocula in IM soil) that were richest in DSE structures and *Alternaria* sequences exhibited a parallel accentuation of negative feedbacks on plant performance (Figs. 4, 5; Table 2), suggesting that these endophytes may mediate synergistic abiotic-biotic feedbacks on plant performance (Waller *et al.*, 2005). DSE such as *Alternaria* are particularly common in boreal and other cooler climates, with effects ranging from highly beneficial (Mandyam & Jumpponen, 2014) to pathogenic (Cheeke *et al.*, 2019). Further work will be necessary to dissect this intriguing biotic interaction across latitudinal and elevational gradients in *M. guttatus*.

Most importantly, annual and perennial *M. guttatus* ecotypes differ not just in field filtering (Fig. 2) but in root-endophyte interactions under controlled conditions, with the DUN perennial exhibiting both higher AMF colonization and apparent taxon-specificity. *Claroideoglomus* (which is the dominant AMF at the DUN field site) predominated in all plants in our mixed inoculum treatment, but the DUN ecotype alone exhibited associations with native-like *Claroideoglomus* in the Glomeraceae-rich IM inoculum (Fig. 6). Although the mechanism is not yet clear and more work will be necessary to confirm this pattern under other conditions, greater filtering by the coastal perennial DUN is consistent with late-successional taxa exhibiting higher specificity in AMF associations (López-García *et al.*, 2014). Perennial life history traits (e.g. higher root biomass or carbon supply) may competitively favor *Claroideoglomus* (Pawlowski *et al.*, 2020; Liu *et al.*, 2020) or DUN may have specific adaptations that maximize colonization by this locally-abundant AMF taxon. This evidence of ecotypic differentiation complements work showing that single mutations to key genes can alter plant-AMF signaling (Pivato *et al.*, 2007) and work documenting species-level differences in plant-AMF associations (Walters *et al.*, 2018; Eck *et al.*, 2019), and provides an avenue into investigating the evolution of host-AMF interactions across diverse *Mimulus*. Along with ecotypic differentiation in negative feedbacks in native biotic-abiotic contexts and differences in AMF colonization under experimental conditions, these results add to the growing evidence that plant genetic variation can shape both directions of feedback with soil microbial communities (Walters *et al.*, 2018; Eck *et al.*, 2019).

## ACKNOWLEDGEMENTS

We thank Gloria Goñi-McAteer and Hanna McIntosh for assistance with field work, Auroralela Bayless and Patrick Demaree for assistance with plant care and performance measures, and Tamara Max of the UM Genomics Core Facility for consultation on the sequencing. Funding for the work was provided by NSF DEB-1457763 to LF, NSF DGE-184053 to MM, Sigma Xi grant-in-aid of research to MM and LF, and MPG Ranch to YL and LB.

## AUTHOR CONTRIBUTIONS

MM, YL, and LF designed the research; MM collected most data; YL quantified fungal root colonization; MM, LF, and LB analyzed data; all authors interpreted data; MM and LF wrote the manuscript, with input and editing from YL and LB.

