## Supporting Information for "Divergence in bidirectional plant-soil feedbacks between montane annual and coastal perennial ecotypes of yellow monkeyflower (*Mimulus guttatus*)"

### New Phytologist Supporting Information

Article acceptance date: Click here to enter a date.

The following Supporting Information is available for this article:

**Fig. S1** Rarefaction curves for field and greenhouse root sequences.

**Table S1** Thermocycler conditions for generation of ITS amplicons and sequencing library preparation.

**Table S2** Field soil nutrient analysis.

**Table S3** Full model for comparison of field fungal communities.

**Table S4** Colonization of field roots.

**Table S5** Performance of DUN and IM ecotypes in virtual reciprocal transplant.

**Table S6** Abundance of fungal sequence variants in inoculum*ecotype combinations.

**Methods S1** Detailed description of inoculum generation.

**Methods S2** Detailed description of sample preparation, amplification, and sequencing for field and greenhouse samples.

**Methods S3** Detailed description of characterization and comparison of RAF communities and FUNGuild analyses.

**Figure S1** Rarefaction curves generated by QIIME II from a) sequenced field root and soil samples and b) greenhouse root samples. For analyses, field and greenhouse samples were rarefied at 1700 and 200 sequences, respectively.

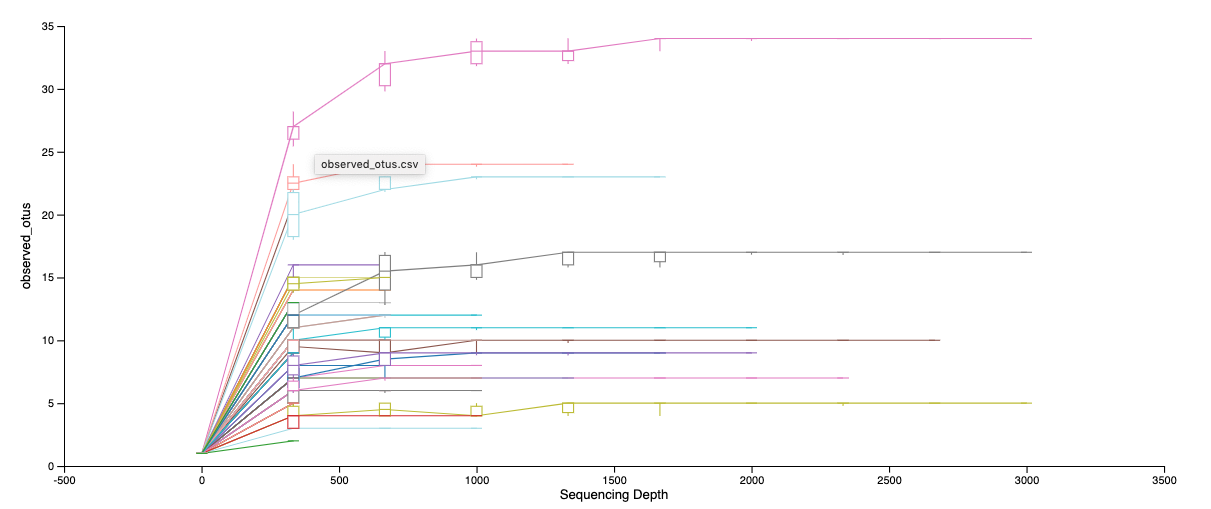

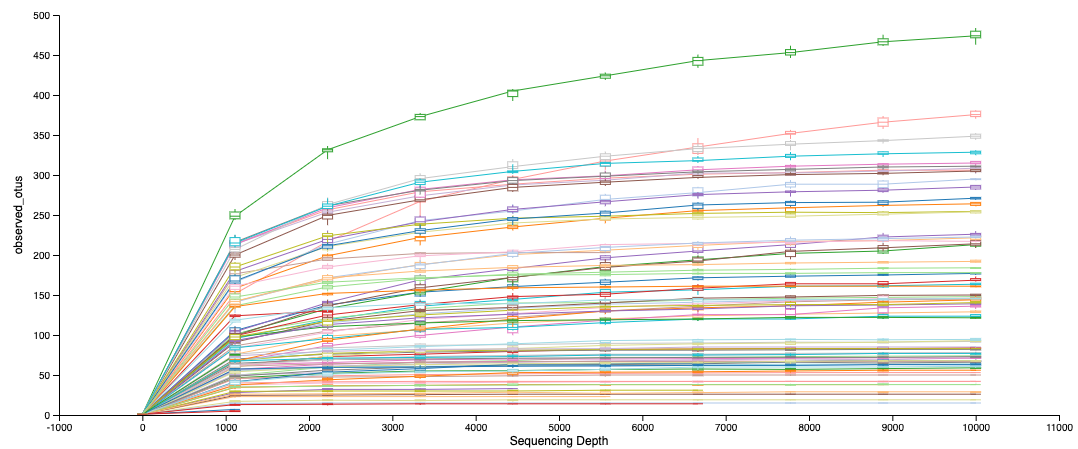

b)

a)

**Table S1** Thermocycler conditions for generation of ITS amplicons and sequencing library preparation.

|  | PCR1 (Amplification) | | PCR2 (Adapter Barcoding ) | |
| --- | --- | --- | --- | --- |
| Step | Temp. (°C) | Time (min) | Temp. (°C) | Time (min) |
| 1 | 94 | 1 | 95 | 1 |
| 2 | 94 | 1 | 95 | 0.5 |
| 3 | 51 | 1 | 60 | 0.5 |
| 4 | 72 | 1 | 68 | 1 |
| 5 | Repeat steps 2-4 29 times | | Repeat steps 2-4 9 times | |
| 6 | 72 | 8 | 68 | 5 |

**Table S2** Field soil nutrient analysis. Bulked soil from each site was analyzed by Ward Laboratories Inc. in Kearney, NE (USA). SOM = soil organic matter.

|  | pH | SOM (%) | N (ppm) | P (ppm) | K (ppm) | S (ppm) | Z (ppm) | Mn (ppm) | Cu (ppm) | Mg (ppm) |
| --- | --- | --- | --- | --- | --- | --- | --- | --- | --- | --- |
| IM | 6.4 | 7.6 | 2.6 | 9 | 182 | 10 | 0.38 | 11.2 | 0.68 | 775 |
| DUN | 6.3 | 0.4 | 0.1 | 7 | 76 | 4 | 0.42 | 1.2 | 0.05 | 52 |

**Table S3** Full model for comparison of field root-associated fungal communities. PERMANOVA of site (DUN, IM), medium (soil, roots), and their interaction as factors explaining Bray-Curtis distances of Hellinger transformed sequence abundances from field samples (N = 17 – 20 per Site*Medium combination).

|  | R^2^ | F-statistic | p-value |
| --- | --- | --- | --- |
| Site | 0.25 | 27.5 | 0.001 |
| Medium | 0.06 | 6.6 | 0.001 |
| Site*Medium | 0.05 | 5.6 | 0.001 |

**Table S4** Colonization of field roots. Percent colonization of field sites by AMF hyphae, AMF vesicles, and non-AMF hyphae. Mean ± standard error reported. P-values are reported in the text.

|  | DUN Site | IM Site |
| --- | --- | --- |
| AMF hyphae | 7.4±2.2 | 11.7±2.0 |
| Non-AMF hyphae | 45.0±4.5 | 54.2±3.6 |

**Table S5** Performance (least squares means ± SE) of DUN and IM M. guttatus ecotypes in a virtual reciprocal transplant into DUN and IM “sites” (home soil + WB biota combinations), with P-values for model effects (2-way ANOVA on ecotype and “site” and their interaction, total N = 34). A Poisson loglinear model for flower number produced the same statistical results, so we present the LSMs and effects from standard ANOVA here.

| Ecotype | DUN | | IM | |  | | |
| --- | --- | --- | --- | --- | --- | --- | --- |
| “Site” (Soil+Biota) | DUN | IM | DUN | IM | ecotype | site | ecotype  *site |
| Days to Flower | 55.8 ± 2.8 | 48.1 ± 2.0 | 33.9± 2.6 | 30.7± 2.1 | <0.0001 | 0.01 | 0.90 |
| Flower Number | 5.8 ± 4.3 | 7.9 ± 4.6 | 31.1 ± 3.9 | 31.0 ± 3.9 | <0.0001 | 0.81 | 0.80 |
| AG Biomass (g) | 0.25 ± 0.01 | 0.47 ± 0.02 | 0.24 ± 0.02 | 0.40 ± 0.02 | 0.25 | 0.01 | 0.12 |
| BG Biomass (g) | 0.21 ± 0.02 | 0.26 ± 0.02 | 0.05 ± 0.02 | 0.11 ± 0.02 | 0.0002 | 0.02 | 0.98 |
| Total Biomass (g) | 0.43 ± 0.05 | 0.61 ± 0.05 | 0.29 ± 0.04 | 0.37 ± 0.04 | 0.0002 | 0.007 | 0.30 |
| Root Mass Ratio | 0.41 ± 0.01 | 0.37 ± 0.01 | 0.18 ± 0.02 | 0.26 ± 0.02 | <0.0001 | 0.50 | 0.02 |

**Table S6** Abundance of fungal sequence variants (assigned to trophic classes and taxa in FUNGuild) found in experimental RAB source*ecotype combinations (sample n in parentheses). Taxa are shown if they were found in >2 samples and had a total of > 20 reads in the full dataset).

|  | | | RAB SOURCE | | | | | | | | | | |
| --- | --- | --- | --- | --- | --- | --- | --- | --- | --- | --- | --- | --- | --- |
|  | | | DUN | | | | DUN+IM | | | | IM | | |
|  |  | | PLANT ECOTYPE | | | | | | | | | | |
| Trophic Class | Taxon | DUN  (12) | | IM  (11) | | DUN  (15) | | IM  (12) | | DUN  (9) | | IM  (10) | |
| Pathotroph | *Plectosphaerella* | 58 | | 226 | | 67 | | 74 | | 0 | | 35 | |
|  | Spizellomycetaceae | 4 | | 19 | | 5 | | 4 | | 0 | | 119 | |
|  | *Venturia* | 19 | | 0 | | 0 | | 2 | | 2 | | 8 | |
| Patho-Saprotroph | Didymellaceae | 113 | | 8 | | 13 | | 9 | | 3 | | 15 | |
|  | Phaeosphaeriaceae | 29 | | 0 | | 38 | | 7 | | 2 | | 6 | |
|  | Pleosporaceae | 88 | | 0 | | 31 | | 89 | | 32 | | 75 | |
| Patho-Sapro-Symbiotroph | *Alternaria* | 6 | | 27 | | 145 | | 345 | | 455 | | 442 | |
|  | Ceratobasidiaceae | 9 | | 33 | | 810 | | 245 | | 142 | | 537 | |
|  | *Fusarium* | 15 | | 19 | | 16 | | 19 | | 0 | | 0 | |
|  | Tremellales | 47 | | 23 | | 1 | | 0 | | 0 | | 0 | |
|  | *Chaetomium* | 0 | | 23 | | 0 | | 0 | | 0 | | 115 | |
| Saprotroph | Orbiliaceae | 59 | | 71 | | 84 | | 32 | | 25 | | 35 | |
|  | Lasiosphaeriaceae | 0 | | 6 | | 4 | | 10 | | 22 | | 0 | |
|  | Hypocreales | 26 | | 156 | | 5 | | 5 | | 0 | | 2 | |
|  | Rutstroemiaceae | 21 | | 0 | | 0 | | 0 | | 160 | | 11 | |
|  | Psathyrellaceae | 1 | | 0 | | 1 | | 22 | | 0 | | 1 | |
| Symbiotroph (all AMF) | *Claroideoglomus* | 195 | | 156 | | 460 | | 259 | | 36 | | 0 | |
|  | Glomeraceae | 0 | | 0 | | 3 | | 11 | | 92 | | 54 | |
|  | *Rhizophagus* | 0 | | 0 | | 0 | | 5 | | 26 | | 11 | |
| Unknown/Other | Pleosporales | | 193 | | 95 | | 170 | | 62 | | 172 | | 113 |
|  | unknown | | 31 | | 172 | | 97 | | 57 | | 135 | | 46 |
|  | *Sebacinale* | | 522 | | 545 | | 542 | | 91 | | 0 | | 18 |
|  | Mucorales | | 0 | | 0 | | 6 | | 20 | | 2 | | 11 |
|  | Chytridiomycota | | 11 | | 53 | | 21 | | 4 | | 0 | | 0 |
|  | *Fragarium* | | 19 | | 0 | | 0 | | 11 | | 0 | | 12 |

**Methods S1** Detailed description of inoculum (RAB source) generation.

The WB inocula includes all fungi, bacteria and other biota, whereas the FB had large-spored fungi removed by filtration {i.e., 20mL soil slurry derived from 50mL of live field inoculum was mixed with water and filtered twice through a 25-micron sieve and filter paper; {Koide:1989jp}}. To concentrate inocula, *M. guttatus* from each field site were grown from seed in native soil mixed with Turface (high-fired calcified clay; PROFILE Products LLC, Buffalo Grove, IL, United States), and sterile sand (1:1:1). Host plants were grown in the greenhouse under cool temperatures and short (12hr) days with light supplementation, low phosphorus fertilization (20mL per 4” pot of 20-2-20 fertilizer at 50ppm N every two weeks) and watering every other day. After three months, we removed plant material, prepared separate WB and FB inocula from each site, and prepared DUN+IM WB and FB inocula by mixing the respective DUN and IM inocula in equal parts. We also generated a generic AMF-rich inoculum (INVAM) and paired FB control using the same methods as for DUN/IM inocula (above). The starting point for these inocula was a mixture of eight AMF species (*Rhizophagus irregularis, R. sinuosum, Gigaspora rosea, G. margarita, Claroideaglomus etunicatum, C. lamellosum, Dentiscutata heterogama* and *Funneliformis mosseae*), sourced from INVAM (https://invam.wvu.edu/) and sub-cultured on *Panicum vulgare* grown for three months in 1:1:1 mix of autoclaved MPG Ranch (Missoula, MT) field soil, sand, and Turface. All inocula were stored for two weeks at room temperature until use. Filtration of inocula eliminated colonization by AMF (mean: WB = 0.16 ± 0.02, n = 42; FB = 0.00 ± 0.04, n = 14) and substantially reduced colonization by non-AMF root endophytes (WB = 0.33 ± 0.03, FB = 0.10 ± 0.06) in experimental roots, suggesting that any differences in plant performance between these treatments may reflect endophyte removal.

**Methods S2** Detailed description of sample preparation, amplification, and sequencing for field and greenhouse samples.

DNA was extracted from 50 -100 mg of greenhouse (fresh) and field (freeze-dried) root samples using a 96-well-format CTAB-chloroform protocol for plant and fungal tissue ([dx.doi.org/10.17504/protocols.io.bgv6jw9e](https://dx.doi.org/10.17504/protocols.io.bgv6jw9e)). For field soil samples, we used PowerLyzer PowerSoil DNA Isolation Kits (MoBio Laboratories, Inc., Solana Beach, CA, USA). Because the small soil volume recommended by the kit (200 mg) resulted in poor DNA yields, we filtered small particles (<60 microns) from a larger volume (15 mL) of field soil and added 200mg of these filtered particles to the kit, then followed the standard kit protocol. From the greenhouse experiment, we sampled roots from the three biotic treatments (DUN, DUN+IM, IM inocula x both plants and soils), including both WB (n=112 before rarefaction, n = 69 after rarefaction) and FB (n=71 before rarefaction, n= 37 after rarefaction) treatments, as well as a small number of plants from the sterile treatment (n=8 before rarefaction, n=4 after rarefaction) and one non-template blank as controls). Each sample was extracted and sequenced once

M**ethods S3** Detailed description of characterization and comparison of RAF communities and FUNGuild analyses.

We used the plugin q2-demux (https://github.com/qiime2/q2-demux) for demultiplexing, trimming forward reads at 240 bases and reverse reads at 200 bases in order to retain the most paired reads. We used q2-dada2 for quality filtering and de-replication using default parameters {Callahan:2016df}. Sequence variants (SVs) were classified using UNITE, an ITS fungal sequence database {Nilsson:2019ka} , and the q2-feature-classifier (https://github.com/qiime2/q2-feature-classifier), a naïve Bayes machine-learning classifier {Bokulich:2018do}. We excluded sequences that did not match UNITE sequences (<85% identity or coverage) to remove non-target DNA. For FUNGuild analyses, we grouped SVs as “mutualists” (arbuscular mycorrhizal, ericoid mycorrhizal, orchid mycorrhizal, or ectomycorrhizal fungi) or as “plant pathogens” (FUNGuild “guild” includes terms “plant pathogen”) , including only SVs with either “probable” or “highly probable” confidence. All other SVs were assigned to “other/unknown”.

**Supplemental Methods References**

**Bokulich NA, Kaehler BD, Rideout JR, Dillon M, Bolyen E, Knight R, Huttley GA, Gregory Caporaso J**. **2018**. Optimizing taxonomic classification of marker-gene amplicon sequences with QIIME 2's q2-feature-classifier plugin. *Microbiome* **6**: 90–17.

**Callahan BJ, McMurdie PJ, Rosen MJ, Han AW, Johnson AJA, Holmes SP**. **2016**. DADA2: High-resolution sample inference from Illumina amplicon data. *Nature Methods* **13**: 581–583.

**Koide RT, Li M**. **1989**. Appropriate controls for vesicular–arbuscular mycorrhiza research. *New Phytologist* **111**: 35–44.

**Nilsson RH, Larsson K-H, Taylor AFS, Bengtsson-Palme J, Jeppesen TS, Schigel D, Kennedy P, Picard K, Glöckner FO, Tedersoo L, *et al.*** **2019**. The UNITE database for molecular identification of fungi: handling dark taxa and parallel taxonomic classifications. *Nucleic Acids Research* **47**: D259–D264.
